# Conserved satellite DNA motif and lack of interstitial telomeric sites in highly rearranged African *Nothobranchius* killifish karyotypes

**DOI:** 10.1101/2023.03.28.534604

**Authors:** Karolína Lukšíková, Tomáš Pavlica, Marie Altmanová, Jana Štundlová, Šárka Pelikánová, Sergey A. Simanovsky, Eugene Yu. Krysanov, Marek Jankásek, Matyáš Hiřman, Martin Reichard, Petr Ráb, Alexandr Sember

## Abstract

Repetitive DNA may have significant impact on genome evolution. African annual killifishes of the genus *Nothobranchius* (Teleostei: Nothobranchiidae), which has adapted to temporary water pools in African savannahs, possess genomes with high repeat content. They are also characterized by rapid karyotype and sex chromosome evolution but the role of genome repeats in these processes remains largely unknown. Here, we analyzed the distribution of telomeric (TTAGGG)*_n_* repeat and Nfu-SatC satellite DNA (isolated formerly from *N. furzeri*) by fluorescence *in situ* hybridization in representatives across the *Nothobranchius* phylogeny (15 species), and with *Fundulosoma thierryi* as an outgroup. All analyzed taxa shared the presence of Nfu-SatC repeat but with diverse organization and distribution on chromosomes (from small clusters scattered genome-wide, to large localized accumulations, or a combined pattern). Nfu-SatC landscape was similar in conspecific populations of *N. guentheri* and *N. melanospilus* but slightly-to-moderately differed between populations of *N. pienaari*, and between closely related *N. kuhntae* and *N. orthonotus*. Inter-individual variability in Nfu-SatC patterns was found in *N. orthonotus* and *N. krysanovi*, including distinct segments present often in heterozygous condition. We revealed mostly no sex-linked patterns of studied repeat’s distribution in any of the sampled species including those with known sex chromosomes. Only in *N. brieni* (having an X_1_X_2_Y multiple sex chromosome system), Nfu-SatC probe covered substantial portion of the Y chromosome, similarly as formerly found in *N. furzeri* and *N. kadleci* (XY sex chromosomes), sister species not closely related to *N. brieni*. All studied species further shared patterns of telomeric FISH, with expected signals at the ends of all chromosomes and no additional interstitial telomeric sites. In summary, we revealed i) the presence of conserved satDNA class in *Nothobranchius* clade (a rare pattern among ray-finned fishes), ii) independent trajectories of *Nothobranchius* sex chromosome diferentiation, with recurrent and convergent accumulation of Nfu-SatC on the Y chromosome in some species, and iii) genus-wide shared propensity to loss of telomeric repeats during the mechanism of interchromosomal rearrangements. Collectively, our findings advance our understanding of genome structure, mechanisms of karyotype reshuffling and sex chromosome differentiation in *Nothobranchius* killifishes from the genus-wide perspective.

## Introduction

Repetitive DNA usually comprises substantial portion of eukaryotic genomes [López-Flores and Garrido-Ramos, 2012; Garrido-Ramos, 2017; Blommaert et al., 2019; Shao et al., 2023]. Entire repetitive content per species’ genome (i.e. repeatome) and its organization on chromosomes feature often species-specific and even population-specific patterns (e.g., [Feliciello et al., 2014; Bracewell et al., 2019; Robledillo et al., 2020]). The repetitive DNA landscape may therefore have a significant impact on the rate of genome evolution, both structurally (i.e. by facilitating chromosome rearrangements; e.g., [Li et al., 2017; Brown and Freudenreich, 2021; Balachandran et al., 2022]) and functionally (by facilitating evolution of genes and their expression; e.g., [Felicielo et al., 2021]). These mechanisms may lead to reproductive isolation between evolving conspecific populations and to subsequent species diversification [Ferree and Barbash, 2009; Lee et al., 2017; Bracewell et al., 2019; Robledillo et al., 2020]. Moreover, repetitive DNA plays diverse important roles in genome organization, maintenance, and function, including regulation of gene expression, proper chromosome segregation during cell divisions, chromatid cohesion, and chromosome end protection [Biscotti et al., 2015; Garrido-Ramos, 2017; Winter et al., 2018; Underwood and Choi, 2019; Louzada et al., 2020].

For instance, specific telomeric tandem repeats being in vertebrates composed of TTAGGG monomeres [Meyne et al., 1990] act in concert with associated protein complexes to protect chromosome ends from being recognized as aberrant double-stranded breaks, and from digestion by nucleases. They also counteract the possibly deleterious effect of genetic material loss during the replication of linear chromosomes [O’Sullivan and Karlseder, 2010; Lazzerini-Denchi and Sfeir, 2016; Vicari et al., 2022].

Satellite DNA (satDNA) is another tandemly-repeated class which is known to have rapid evolution rate resulting in high genomic diversity both qualitatively (i.e. monomer nucleotide sequence, size and complexity) and quantitatively (amount and genomic organization) [Plohl et al., 2012; Garrido-Ramos, 2017; Thakur et al., 2021]. These sequences represent major component of densely packed constitutive heterochromatin [Plohl et al., 2012; Garrido-Ramos, 2017] and may also play important role in centromere organization and function [Melters et al., 2013; Hartley and O’Neill, 2019; Talbert and Henikoff, 2020]. SatDNA classes whose entire genomic collection is referred to as satellitome [Ruiz-Ruano et al., 2016] usually form long tracts of tandem repeats which are being intraspecifically homogenized by mechanisms of molecular drive, leading to so-called concerted evolution [Plohl et al., 2012]. They can also show non-clustered organization or form smaller clusters dispersed across the genome, even in euchromatic parts [Ruiz-Ruano et al., 2016; Crepaldi and Parise-Maltempi, 2020; Šatović-Vukšić and Plohl, 2023]. SatDNA evolves under relaxed purifying selection unless gaining a role in cellular function [Garrido-Ramos, 2017; Lower et al., 2018]. While new satDNAs may emerge and some resident ones completely disappear from the particular genome through the evolutionary time, a collection of shared satDNA motifs is believed to exist in related species, following the concept of so-called “library” hypothesis [Fry and Salser, 1977]. What then differs inter-specifically, or between populations of the same species, is the degree of expansion or contraction of particular repeat motifs along with their dissemination in new genomic regions and further sequence evolution. A variety of underlying mechanisms for these processes have been described (e.g., [Ruiz-Ruano et al., 2016; Camacho et al., 2022; Šatović-Vukšić and Plohl, 2023]).

In teleost fishes, similarly to other organisms, repetitive DNA classes serve as chromosomal landmarks for tracking lineage-specific genome, karyotype and sex chromosome dynamics, and for cytotaxonomic purposes (e.g., [Bellafronte et al., 2005; Schemberger et al., 2011; Cioffi and Bertollo, 2012; Glugoski et al., 2020; Goes et al., 2020; Yano et al., 2021]). Most widely used repeats have been so far ribosomal DNA (rDNA) clusters and telomeric sequences, due to their deep sequence conservation and mostly easy visualization by fluorescence *in situ* hybridization (FISH) [Cioffi and Bertollo, 2012; Ocalewicz, 2013; Sochorová et al., 2018; Vicari et al., 2022]. Developments in genomics and in bioinformatic pipelines during the last decade greatly boosted the analysis of highly variable and formerly hardly tractable repeats, such as mobile elements and satDNA (e.g., [Ruiz-Ruano et al., 2016; Garrido-Ramos, 2017; Lower et al., 2018; Novák et al., 2020; Vondrak et al., 2020; Šatović-Vukšić and Plohl, 2023]). Satellitome analyses in teleosts helped, for instance, in characterization of fish interspecific hybrids [Marta et al., 2020], supernumerary (B) chromosomes [Serrano-Freitas et al., 2020] and sex chromosomes [Schemberger et al., 2019; Utsunomia et al., 2019; Crepaldi and Parise-Maltempi, 2020; Kretschmer et al., 2022]. Nonetheless, the satellitome research has so far centred mainly on Neotropical characiform lineages (e.g., [Utsunomia et al., 2019; dos Santos et al., 2021; Kretschmer et al., 2022; Goes et al., 2023]). Therefore, to achieve a more general understanding of teleost satellitome and its impact on genome evolution, a broader spectrum of non-related fish groups should be surveyed. African killifish genus *Nothobranchius*Peters, 1868 (Aplocheiloidei: Nothobranchiidae) comprises more than 90 recognized species [Nagy and Watters, 2021; Fricke et al., 2022] partitioned into seven main evolutionary clades [van der Merwe et al., 2021]. These small-bodied fishes with marked sexual dimorphism and dichromatism (males are more colorful and larger than females) [Wildekamp, 2004; Berois et al., 2016] live in seasonal wetlands in south-east African savannah regions [Wildekamp, 2004]. Due to the transient nature of this environment, *Nothobranchius* spp. evolved key adaptations to endure the dry period, such as desiccation-resistant eggs and fast life cycle to accomplish reproduction before water pools dry out [Cellerino et al., 2016; Furness, 2016]. The genus includes the turquoise killifish (*N. furzeri* Jubb, 1971), a model for ageing research as its wet part of the life cycle represents the shortest lifespan among vertebrates recorded in captivity [Cellerino et al., 2016; Hu and Brunet, 2018].

All *Nothobranchius* spp. live in small, geographically isolated populations and are thus subject to genetic drift, including bottlenecks and founder effects [Cui et al., 2019], which facilitates accumulation of genetic polymorphisms [King, 1993; Nonaka et al., 2019]. This effect is further enhanced by assortative mating and inbreeding [Bartáková et al., 2013; Berois et al., 2016; Willemsen et al., 2020]. These processes and the ability to disperse and colonize new water ponds during major floods in rainy season shaped considerably the genome evolution and diversification in these fishes [Bartáková et al., 2013, 2015; Dorn et al., 2014; van der Merwe et al., 2021].

*Nothobranchius* genomes harbour a high amount of repetitive DNA (about 60–80 %; [Reichwald et al., 2009, 2015; Cui et al., 2019; Štundlová et al., 2022]) which, together with their specific demographic characteristics, might have contributed to remarkable karyotype variability reported thus far among 73 studied *Nothobranchius* representatives [Krysanov et al., 2016, 2023; Krysanov and Demidova, 2018]. Numerous structural rearrangements (mainly fusions, translocations and fissions) resulted in wide range of diploid chromosome numbers (2n = 16–50) in several lineages [Krysanov and Demidova, 2018], while other ones remained more conservative in this sense [Krysanov et al., 2023].

Furthermore, certain *Nothobranchius* spp. have been found to possess sex chromosomes. A standard ♀XX/♂XY sex chromosome system was originally described by genomic approach in *N. furzeri* [Reichwald et al., 2015] and later by cytogenomics in its sister species *N. kadleci* [Štundlová et al., 2022]. Six other *Nothobranchius* representatives as well as the sister species of this genus, *Fundulosoma thierryi*, evolved multiple sex chromosomes of the ♀X_1_X_1_X_2_X_2_/♂X_1_X_2_Y type, most probably after a Y-autosome fusion (meaning that X_1_ denotes the original X chromosome and X_2_ represents the unfused autosomal homolog) [Ewulonu et al., 1985; Krysanov et al., 2016; Krysanov and Demidova 2018]. Except for demonstrably shared origin of XY sex chromosome system in *N. furzeri* and *N. kadleci* [Štundlová et al., 2022], the relationships between the remaining sex chromosome systems are still unknown. Representatives carrying X_1_X_2_Y sex chromosomes, namely *N. guentheri*, *N. brieni*, *N. ditte*, *N. janpapi*, *N. lourensi* and *Nothobranchius* sp. Kasenga, occupy different positions in the *Nothobranchius* phylogeny, nested with many species with unknown sex determination mode [Krysanov and Demidova, 2018; van der Merwe et al., 2021]. It is therefore presumed that these systems had recurrent and independent origins [Krysanov and Demidova, 2018] but a solid evidence is still missing. Nothing is further known about the genetic content and degree of differentiation of these sex chromosomes except for *N. guentheri* (and outgroup *F. thierryi*) where banding techniques and physical mapping of certain repeats did not show considerable repeat accumulation on neo-Y [Voleníková et al., 2023].

Štundlová et al. [2022] analyzed repeatome of *N. furzeri* and *N. kadleci* by RepeatExplorer2 pipeline [Novák et al., 2020] and found the following shared characteristics: two most abundant repeats (Nfu-SatA and Nfu-SatB) occupied (peri)centromeric regions of all chromosomes, while another repeat, Nfu-SatC, was dispersed across the chromosome complement but formed a prominent specific accumulation on the heteromorphic Y chromosome.

In our complementary study [Voleníková et al., 2023], we probed Nfu-SatA, Nfu-SatB and other satDNA motifs on chromosomes of species of Southern and Coastal clade. We revealed fast turnover of (peri)centromeric satDNA repeats and possible indications of the action of meiotic drive.

In the present study, we aimed to track footprints of chromosome rearrangements and the presence of differentiated sex chromosomes. We therefore investigated the patterns of distribution of Nfu-SatC satDNA and telomeric repeats in representative sampling encompassing 15 *Nothobranchius* species from four evolutionary lineages (Southern, Ocellatus, Coastal and Kalahari clade) and *F. thierryi* as an outgroup. While none of the other so far mapped repeats [Voleníková et al., 2023] indicated the existence of yet uncharacterized sex chromosomes in surveyed species, the mapping of Nfu-SatC may be particularly promising in this sense thanks to its assumed ability to accumulate preferably in regions of suppressed recombination [Štundlová et al., 2022]. In the three out of five studied species with X_1_X_2_Y sex chromosomes, we further assessed the distribution of constitutive heterochromatin, to complement already known patterns studied by Voleníková et al. [2023]. We found a conserved presence of Nfu-SatC across the *Nothobranchius* phylogeny and the outgroup species, but sex chromosome-specific accumulation in *N. brieni* only. Telomeric FISH further suggested commonalities in general trajectory of structural rearrangements in *Nothobranchius* lineage.

## Materials and Methods

### Fish Species Sampling

We analyzed 63 individuals belonging to 15 *Nothobranchius* killifish species (18 populations) from four phylogenetic clades, and the outgroup species *Fundulosoma thierryi*. The studied individuals from *N. orthonotus*, *N. kuhntae*, *N. pienaari*, *N. rachovii*, *N. eggersi*, *N. rubripinnis* and *N. melanospilus* were sampled from laboratory populations recently derived from wild-caught individuals and were previously identified based on morphology and the phylogenetic analysis of mitochondrial and nuclear markers (see [Bartáková et al., 2015; Blažek et al., 2017; Reichard et al., 2022a, b]). The remaining species were obtained from specialists and experienced hobby breeders who keep strictly population-specific lineages derived from original imports. In this case, the species identity was confirmed on the basis of key morphological characters [Wildekamp, 1996, 2004; Watters et al., 2008, 2020; Nagy, 2018]). The sampling is summarized in Figure 1. Detailed information is provided in Supplementary Table 1.

**Figure 1.**
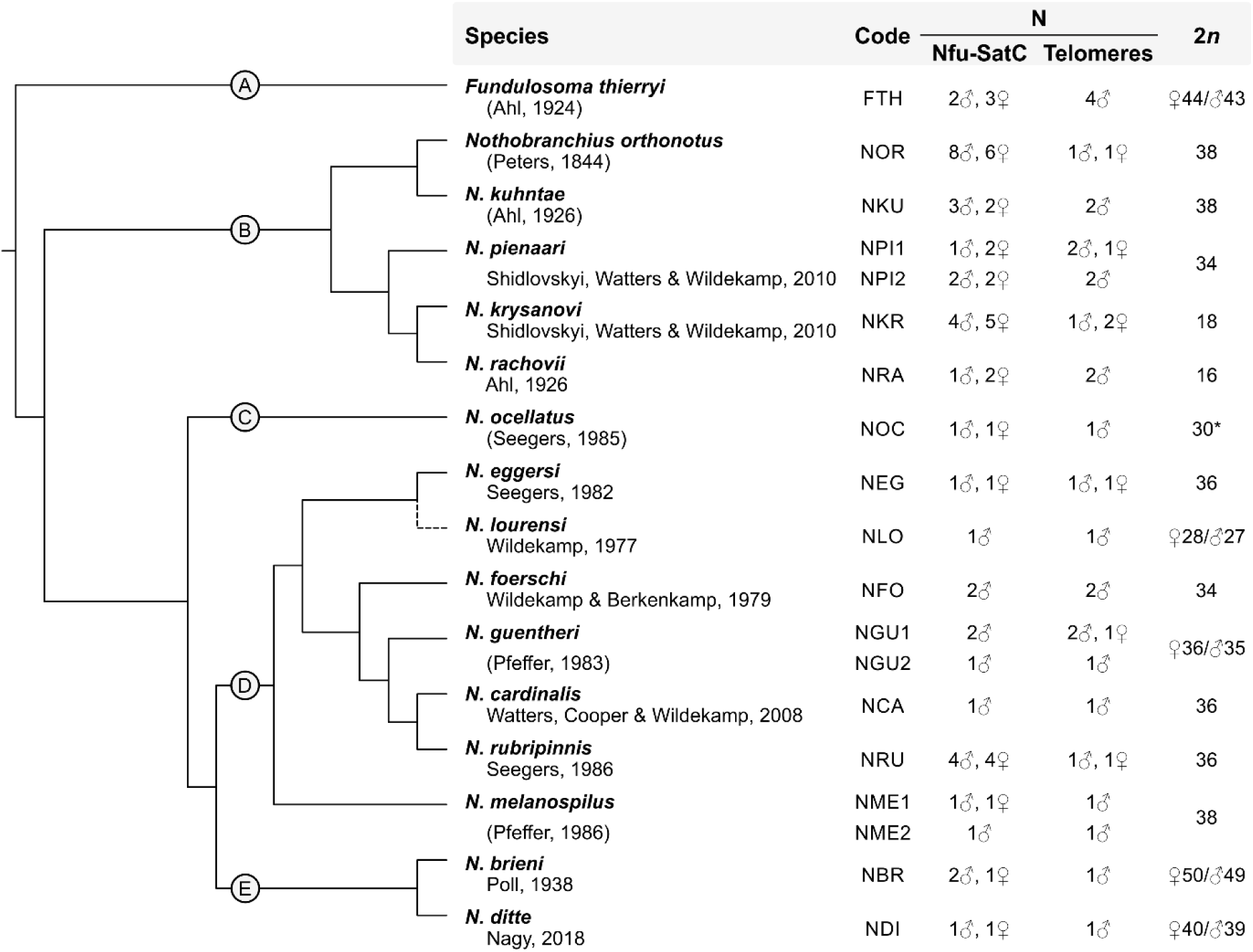
List of *Nothobranchius* killifish species and populations used in this study, with assignment to their phylogenetic positions (following van der Merwe et al. [2021]) and sample sizes (N) per method, and with anchored diploid chromosome numbers (2n) inferred from Krysanov and Demidova [2018]. Asterisk points on a single case where 2n is discordant with data revealed by us in the present study (for details, see main text). The exact placement of *N. lourensi* in the phylogeny has not been determined so far; for the purposes of this study, it was roughly estimated based on morphological characters and geographic distribution [M. Reichard, pers. commun.]. The phylogenetic position of *N. lourensi* is therefore demarcated by a dashed line. Additional details about the sampled *Nothobranchius* populations are provided in online suppl. Table 1. clades: A = Fundulosoma; B = Southern; C = Ocellatus; D = Coastal; E = Kalahari.

### Conventional Cytogenetics and Chromosome Banding

Mitotic chromosome spreads were obtained either from regenerating caudal fin tissue or from cephalic kidney. In the former, we followed Völker and Ráb [2015], with the modifications by Sember et al. [2015], and with an altered time of fin regeneration (one-to-two weeks). In the latter, the kidney-derived chromosome preparations, we followed either Ráb and Roth [1988], or Kligerman and Bloom [1977]. The latter protocol was modified according to Krysanov and Demidova [2018]. The quality of chromosomal spreading on slides was improved following Bertollo et al. [2015]. The amount and quality of chromosome spreads was inspected under phase-contrast optics and suitable slides were dehydrated in an ethanol series (70%, 80% and 96%, 3 min each) before storage in a freezer (−20 °C).

In *N. brieni*, *N. ditte* and *N. lourensi*, i.e. three species with X_1_X_2_Y sex chromosome system, constitutive heterochromatin distribution was assessed by C-banding [Haaf and Schmid, 1984], using 4′,6-diamidino-2-phenolindole (DAPI) (1.5 μg/mL in anti-fade; Cambio, Cambridge, UK) counterstaining. The only protocol modification was done in *N. brieni*: due to observed higher sensitivity of chromatin to barium hydroxide treatment, the incubation time of this step was shortened from 3 to 2.5 min. The C-banding profiles of *F. thierryi*, *N. guentheri* and majority of other herein studied species were subject of a complementary study [Voleníková et al., 2023].

### Nfu-SatC FISH Probe Preparation

We used cloned fragment of Nfu-SatC prepared and verified in the previous study [Štundlová et al., 2022]. This specific clone (GenBank ID: OM542184) contains two adjacent fragments of Nfu-SatC motif which together encompass the entire Nfu-SatC monomere length (634 bp according to RepeatExplorer2 analysis; Štundlová et al. [2022]). The Nfu-SatC fragments form a continuous array, uninterrupted by other sequences in between. The entire plasmid was used as a template for nick translation using a Cy3 NT Labeling Kit (Jena Bioscience, Jena, Germany). For the final probe mixture preparation, 250–500 ng of labeled plasmid and 12.5–25 μg of sonicated salmon sperm DNA (Sigma-Aldrich) were used per slide. The final hybridization mixtures for each slide (15 μL) were prepared according to Sember et al. [2015].

### FISH Analyses

FISH was performed using a combination of two previously published protocols. Specifically, slide pre-treatment, probe/chromosomes denaturation and hybridization followed Sember et al. [2015], while post-hybridization washing was done according to Yano et al. [2017]. The slight protocol modifications were described in Štundlová et al. [2022]. Briefly, following the standard pre-treatment steps, chromosome denaturation was done in 75% formamide in 2× SSC (pH 7.0) (Sigma-Aldrich) at 72 °C, for 3 min. The hybridization mixture which was denatured for 6 min at 86 °C was then spotted onto the slide and the hybridization took place overnight (12–17 h) in a moist chamber at 37 °C. Subsequently, a series of post-hybridization washes was done as follows: two times in 1× SSC (pH 7.0) (65 °C, 5 min each), once in 4× SSC in 0.01% Tween 20 (42 °C, 5 min), and once in 1× PBS (1 min). Slides were then passed through an ethanol series and mounted in antifade containing 1.5 µg/mL DAPI (Cambio, Cambridge, United Kingdom).

Regarding chromosomal mapping of telomeric (TTAGGG)*_n_* repeats, we used a commercial telomere PNA probe directly labeled with Cy3 (DAKO, Glostrup, Denmark). A single modification to the manufacturer’s protocol was the prolonged hybridization time (1.5 h).

### Microscopic Analyses and Image Processing

Images from all cytogenetic methods were captured under immersion objective 100× using BX53 Olympus microscope equipped with appropriate fluorescence filter set, and coupled with a black and white CCD camera (DP30W Olympus). Images were acquired for each fluorescent dye separately using DP Manager imaging software (Olympus). The same software was used to superimpose the digital images with the pseudocolors (blue or red for DAPI, red or green for Cy3; for specific colour coding, see figure legends). Composite images were optimized and arranged using Adobe Photoshop, version CS6.

At least 20 chromosome spreads per individual in each method were analyzed, some of them sequentially. The chromosome morphology description followed Levan et al. [1964], but modified the classification as m – metacentric, sm – submetacentric (biarmed chromosomes), st – subtelocentric, and a – acrocentric (monoarmed chromosomes).

## Results

### Karyotype Characteristics and Distribution of the Constitutive Heterochromatin

Cytogenetic characteristics (2n, karyotype structure) of individuals from all studied killifish species mostly agreed with previous reports [Reichwald et al., 2009, 2015; Krysanov and Demidova, 2018] (Figures 1–3). In *N. pienaari*, *N. guentheri* and *N. melanospilus*, 2n was consistent also at the inter-population level. Only in *N. ocellatus*, both herein studied individuals displayed 2n = 32 composed of monoarmed chromosomes only, while Krysanov and Demidova [2018] reported 2n = 30 for their two studied individuals, where the largest chromosome pair was metacentric. It was later found that individuals studied by Krysanov and Demidova [2018] belong to a newly established species *N. matanduensis* [Watters et al., 2020; S. Simanovsky, pers. commun].

Furthermore, in line with the previous reports [Krysanov et al., 2016; Krysanov and Demidova, 2018], *Fundulosoma thierryi*, *N. brieni*, *N. guentheri*, *N. ditte* and *N. lourensi* possessed male-heterogametic X_1_X_2_Y multiple sex chromosome system, manifested by consistent differences in 2n between males and females (males had one chromosome less than females). Neo-Y chromosomes could be unambiguously identified based on their size and morphology in males of *N. guentheri* and *N. lourensi* (large unpaired submetacentric and subtelocentric chromosome, respectively), or based on hybridization patterns in *N. brieni* (see below).

C-banding in *N. brieni*, *N. ditte* and *N. lourensi* revealed diverse patterns of constitutive heterochromatin distribution (Supplementary Figure 1). *N. brieni* featured high amount of constitutive heterochromatin concentrated in about one-third of the chromosome set. In several chromosomes, heterochromatin blocks were particularly prominent and some of them covered most of the chromosome length. A heterochromatin block stained less intensely by C-banding occupied also short (p) arms of submetacentric neo-Y chromosome. Intriguingly, the heterochromatin blocks only rarely overlapped with Nfu-SatC landscape in this genome (Supplementary Figure 1A–F). In *N. ditte* with relatively low constitutive heterochromatin content, clear heterochromatin blocks (of variable sizes but mostly thick) were confined to (peri)centromeric regions and p-arms of about half portion of the chromosome complement (Supplementary Figure 1G). In *N. lourensi*, C-bands were observed in the (peri)centromere of each chromosome and additional terminal or interstitial signals were also apparent. Particularly long (q) arms of neo-Y chromosome possessed two interstitial heterochromatin blocks (Supplementary Figure 1H).

### Repetitive DNA Patterns as Revealed by FISH

FISH with Nfu-SatC probe revealed consistent signals in all analysed species and populations, including the outgroup *F. thierryi* (Figure 2). The amount and distribution patterns of this satDNA were profoundly different among species. They further slightly differed between the two populations of *N. pienaari* while they were very similar between conspecific populations in *N. guentheri* and *N. melanospilus*. The three following general patterns of repeat arrays organization could be identified: i) many small clusters dispersed across chromosomes, ii) larger clusters restricted to particular chromosomal regions, mostly around centromeres, iii) mixed pattern. Exclusively dispersed organization was recorded in *N. cardinalis*, *N. foerschi*, *N. melanospilus* (Coastal clade) and *N. ditte* (Kalahari clade). A mixed pattern of small dispersed signals mixed with larger localized accumulations was revealed in *F. thierryi* (outgroup), in *N. orthonotus*, *N. kuhntae*, *N. pienaari* (Southern clade) and *N. rubripinnis* (Coastal clade). In *N. pienaari*, the Nfu-SatC landscape in population NPI2 differed from that of NPI1 in the way that the chromosome set largely lacked the small dispersed clusters. Remaining seven studied species possessed larger localized accumulations, mostly in (peri)centromeric regions of many (but never all) chromosomes in the complement. In *N. rachovii*, for instance, Nfu-SatC accumulation was confined to centromeres of six out of 16 chromosomes; some of the clusters were remarkably large. In *N. orthonotus* and *N. krysanovi*, polymorphic blocks of Nfu-SatC accumulations were observed, often in heterozygous condition (Figure 2B, C, H and Supplementary Figure 2). While in *N. krysanovi* we observed a single polymorphic site (large accumulation present on both or just one homolog of a submetacentric pair), in *N. orthonotus* the variablity was present on several chromosomes and it also sometimes resulted in heteromorphism in size and morphology between homologs (we focused on a more detailed analysis of three chromosomal pairs with the most notable polymorphic regions; Supplementary Figure 2). Interestingly, this type of variability was not present in the closest related species *N. kuhntae*. The most prominent Nfu-SatC accumulation in *N. orthonotus* was observed on a pair of large submetacentric chromosomes where it displayed a varied degree of array amplification. All these polymorphisms were sex-unrelated, i.e. present in individuals of both sexes. The only sex-specific accumulation was revealed in *N. brieni*, where Nfu-SatC clusters covered almost the entire length of the neo-Y chromosome in males (Figure 2T).

**Figure 2.**
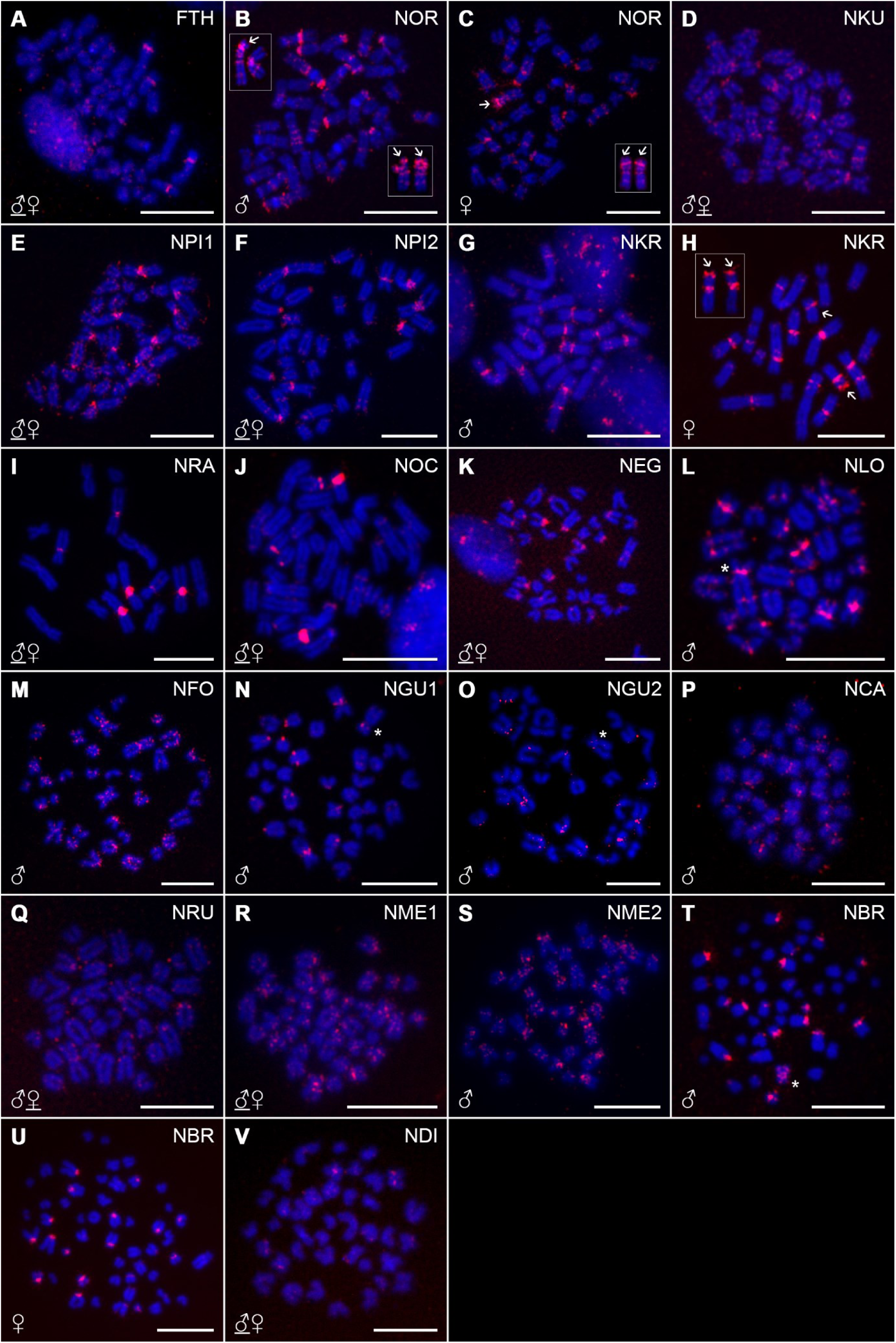
Mitotic metaphases of *Nothobranchius* spp. after single-colour FISH with Nfu-SatC probe (red signals). Chromosomes were counterstained with DAPI (blue). Sex of the studied individuals is indicated and eventually underlined where both sexes (if studied) presented the same distribution pattern (the presented metaphase belongs to individual of underlined sex). In species with high signal pattern variability at certain positions (*N. orthonotus* – B, C; *N. krysanovi* – G, H), and with sex-specific pattern (*N. brieni* – T, U), metaphases of both sexes are presented. Chromosomes with a polymorphic Nfu-SatC patterns are framed; arrows point to segments of high repeat accumulation. More detailed analysis of *N. orthonotus* – linked signal pattern polymorphisms are provided in Supplementary Figure 2. Asterisks correspond to detectable Y chromosomes in species with X_1_X_2_Y sex chromosome system. Note remarkable Nfu-SatC accumulation on the Y chromosome of *N. brieni* (T). Species order reflects their phylogenetic relationships ([van der Merwe et al., 2021] and Figure1). Species coding follows Figure 1. Scale bar = 10 μm.

By mapping the distribution of the telomeric sequences we revealed the standard pattern with signals at the ends of all chromosomes, as expected. Non-telomeric signals have not been retrieved in any species under study, including those with multiple X_1_X_2_Y sex chromosomes (Figure 3). A remarkable signal pattern was revealed in *N. cardinalis* where the telomeric repeat probe generated strong signals in the terminal regions of the p-arms of all acrocentric chromosomes, while the signals at the ends of the q-arms of these chromosomes were notably weaker. Finally, the telomeric probe was unable to detect telomeric repeats in the only pair of large metacentric chromosomes in the *N. cardinalis* karyotype (Figure 3N), as verified also under enhanced exposition times.

**Figure 3.**
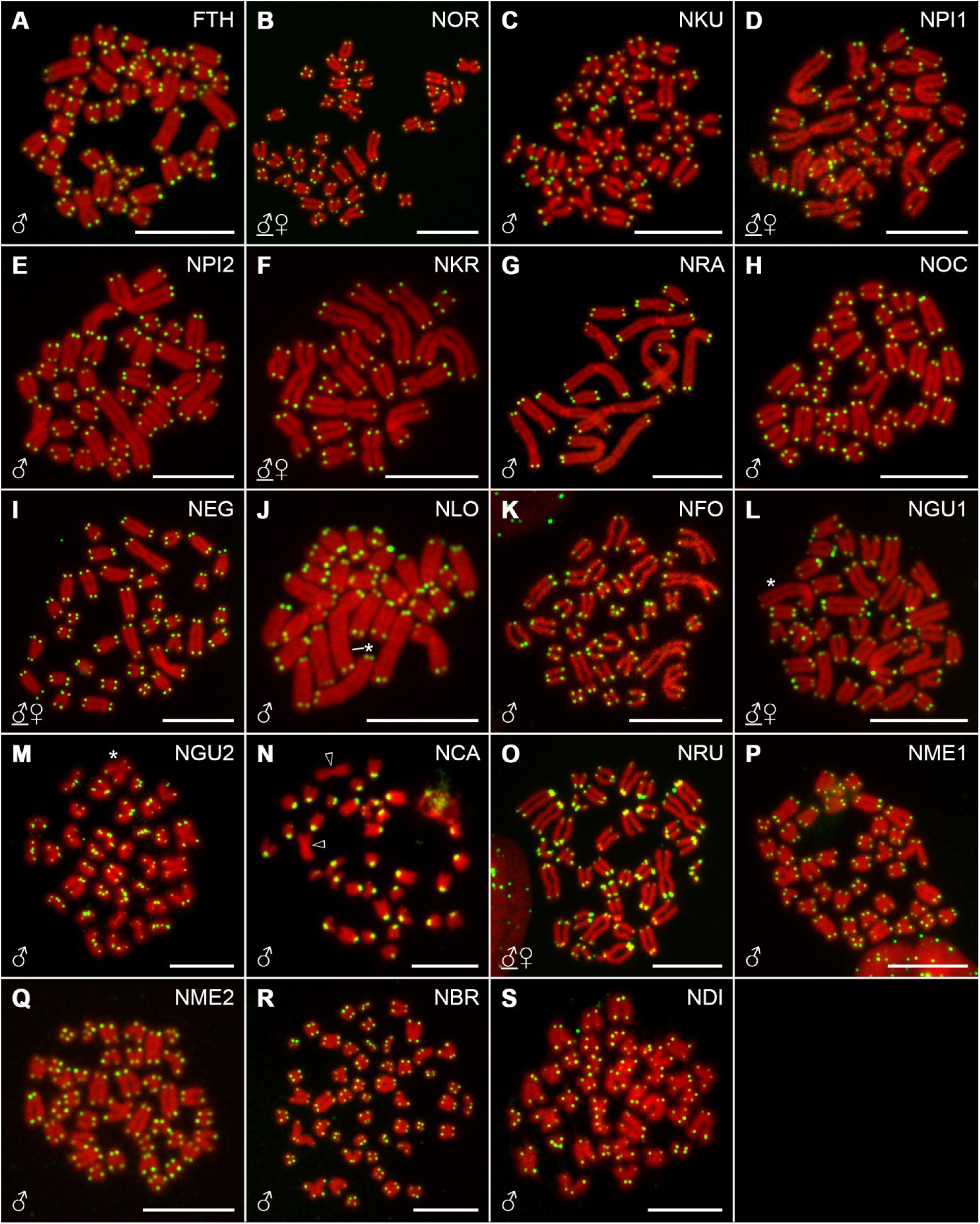
Mitotic metaphases of *Nothobranchius* spp. after PNA-FISH with telomeric probe. For better contrast, pictures were pseudocolored in green (telomeric repeat probe) and red (DAPI counterstaining). Sex of the studied individuals is indicated and eventually underlined where both sexes were studied, to demarcate the origin of the metaphase spread presented. Asterisks correspond to detectable Y chromosomes in species with X_1_X_2_Y sex chromosome system. Note lack of interstitial telomeric sites (ITSs) in the entire sampling, including large Y chromosomes. In *N. cardinalis* (N) empty arrowheads point to the pair of large metacentric chromosomes which lack detectable telomeric signals. Species order reflects their phylogenetic relationships ([van der Merwe et al., 2021] and Figure 1). Species coding follows Figure 1. Scale bar = 10 μm

## Discussion

In the present study, we assessed the distribution patterns of two selected repetitive sequences which had a potential to provide important landmarks for interpretation of karyotype changes and sex chromosome evolution in *Nothobranchius* killifishes. We performed the analysis on the representative sampling of 15 species from different *Nothobranchius* clades. At the same time, our study for the first time indicates divergent processes of multiple sex chromosome differentiation among their species.

For comparative mapping, we chose a telomeric repeat motif which might be instrumental in revealing breakpoints of previous rearrangements [Meyne et al., 1990; Ocalewicz, 2013] and one of the three satellite DNA repeats identified and chromosomally mapped in *N. furzeri* and *N. kadleci* genomes, namely Nfu-SatC [Reichwald et al., 2015; Štundlová et al., 2022]. Nfu-SatC satDNA forms tandem arrays of variable lengths scattered across *N. furzeri* and *N. kadleci* chromosomes but with a remarkable enrichment on the Y sex chromosome of these two species [Štundlová et al., 2022].

Intriguingly, we found detectable clusters of Nfu-SatC in all species under study including the outgroup species, *F. thierryi*, which diverged from the lineage leading to *Nothobranchius* approx. 15 million years ago (MYA) [van der Merwe et al., 2021]. Given that our analysis encompasses four *Nothobranchius* evolutionary lineages including late-branching Kalahari clade, we may assume that Nfu-SatC repeat is highly likely conserved genus-wide and is hence at least 15 MY old. Cases of conserved satDNA in ray-finned fishes are very rare: PstI in sturgeons (more than 100 MY old; [Robles et al., 2004]) and CharSat01 in Characoidei (78– 140 MY old; [dos Santos et al., 2021]). Overwhelming majority of satDNA motifs has otherwise fast turnover [Garrido-Ramos, 2017].

The long-time evolutionary preservation of certain satDNA in a given lineage might be driven by selection, after the repeat motif acquires a specific role in cellular function [Melters et al., 2013; Garrido-Ramos, 2017; Louzada et al., 2020; Feliciello et al., 2021]. Our present data, however, show highly varied patterns of Nfu-SatC distribution in genomes of different *Nothobranchius* species, therefore, the above explanation seems not to apply here. More probably, Nfu-SatC has been so far maintained in *Nothobranchius* genomes stochastically, by neutral processes [Camacho et al., 2022]. This view is supported by the fact that Nfu-SatC patterns largely do not follow phylogenetic relationships. The possible underlying reasons for stochastic Nfu-SatC maintenance might be that i) satDNAs with longer motifs (such as Nfu-SatC) degenerate more slowly at sequence level over time and that ii) dispersion into many smaller clusters across the genome reduces the efficiency of concerted evolution [Camacho et al., 2022; Šatović-Vukšić and Plohl, 2023]. If this reasoning is correct, then Nfu-SatC motif would be more likely preserved throughout the evolutionary time as its small dispersed clusters are being less impacted by concerted evolution. An opposite trend has been recently proposed for serrasalmine fishes [Goes et al., 2023], where the authors suggested that more clustered satDNA repeat organization leads to higher intraspecific homogeneity but higher interspecific repeat differentiation over time.

Transposable elements (TEs) might have helped spreading Nfu-SatC to new genomic locations [Vondrak et al., 2020; Zeljko et al., 2020; Šatović-Vukšić and Plohl, 2023]. Indeed, several retrotransposons have been found among the set of highly abundant repeats in *N. furzeri* and *N. kadleci* genomes [Štundlová et al., 2022]. While they have not been found bioinformatically to form composed repeat clusters with Nfu-SatC [Štundlová et al., 2022], preliminary cytogenetic observations [Sember A., unpublished data] show that some of them display similar dispersed organization across the chromosome complement in *N. furzeri* and *N. kadleci* as does Nfu-SatC repeat. Experiments with simultaneous mapping of Nfu-SatC with these repeats are needed to further assess the effect of TEs on Nfu-SatC dynamics.

It seems that, given its ubiquity, Nfu-SatC stochastically populated regions of reduced/abolished recombination due to relaxed purifying selection (cf. [Stephan, 1986; Charlesworth et al., 1994; Garrido-Ramos, 2017; Kent et al., 2017]). Nfu-SatC hence occupies particularly i) (peri)centromeres of some chromosomes (best seen in *N. rachovii*), ii) regions which might coincide with supposed breakpoint junctions (on the large Y chromosome of *N. lourensi*) and iii) regions of advanced sex chromosome differentiation (Y chromosome of *N. brieni*; discussed further below). Unexpectedly, the specific case of *N. brieni* demonstrates that Nfu-SatC distribution profile may be largely uncorrelated with the distribution of constitutive heterochromatin which typically occupies regions of suppressed recombination (Supplementary Figure 1A–F). Furthermore, particularly interesting is the situation in *N. orthonotus* and *N. krysanovi*, where Nfu-SatC showed polymorphic sites between members of the same chromosome pair. The most remarkable pattern was found in *N. orthonotus* where several chromosomes showed polymorphic sites which might have contributed also to the size differences observed between homologs Supplementary Figure 2). Some of the polymorphic accumulations of Nfu-SatC especially on a pair of large metacentric chromosomes, though sex-unrelated, resemble the pattern observed on the Y chromosomes of *N. furzeri* and *N. kadleci*. Two outcomes might result from this observation. Firstly, the polymorphic region may represent region of suppressed/reduced recombination after some type of structural rearrangement such as inversion. Location of Nfu-SatC block on either p- or q-arms of the chromosome in question (Supplementary Figure 3) might favor this view as particularly pericentric inversion may change the ratio of chromosome arms. Secondly, given the random occurrence of heterozygous condition in both males and females, this chromosome pair with Nfu-SatC accumulations highly likely does not represent sex chromosomes. This inferrence corroborates our previous findings [Štundlová et al., 2022] suggesting that XY sex chromosome system evolved in a common ancestor of *N. furzeri* and *N. kadleci* and hence is not shared with the closest related species *N. orthonotus*. It will be, however, interesting to find out in the future studies whether this polymorphic chromosome pair in *N. orthonotus* involves linkage group representing XY sex chromosomes in *N. furzeri* and *N. kadleci* given that regions with low recombination rates (such as polymorphic structural variants) may be generally predisposed to evolve sex-linked regions [Charlesworth, 2019]. An indirect evidence suggesting that *N. orthonotus* determines sex by different mechanism (i.e. different sex chromosomes or by other means) could be that heterospecific mating between male *N. furzeri* and female *N. orthonotus* produces progeny with reduced viability [Polačik and Reichard, 2011]; however, the underlying reasons for asymmetric hybridization may be more complex (cf. [MacPherson et al., 2023]).

Together with Neotropical *Harttia* catfishes [Deon et al., 2020; Sassi et al., 2020], *Nothobranchius* killifishes show the highest incidence of multiple sex chromosomes at the genus level among teleosts discovered to date [Sember et al., 2021]. It is therefore highly informative lineage for investigating underlying evolutionary forces that drive transitions from standard sex chromosomes (or other forms of sex determination) to multiple sex chromosome systems (cf. [Sember et al., 2021]) and the evolutionary aspects of sex chromosome turnover in general [Vicoso, 2019]. Our present analysis utilized mapping of Nfu-SatC as it proved to be informative in identifying differentiation on XY sex chromosomes of *N. furzeri* and *N. kadleci* [Štundlová et al., 2022]. We, however, did not reveal any demonstrably sex-linked Nfu-SatC accumulation in our sampling except for its massive accumulation on the neo-Y chromosome of *N. brieni*. This means that i) known multiple sex chromosomes in *Nothobranchius* follow different trajectories of differentiation and that ii) other species either possess homomorphic sex chromosomes at a very early stage of differentiation, or accumulate yet unstudied repeats [Voleníková et al., 2023; this study], or govern sex determination by other mechanisms (cf. [Shen and Wang, 2019; Lichilín et al., 2023; Schartl et al., 2023]). Nfu-SatC is therefore not an universal marker for sex chromosome identification in *Nothobranchius* clade. Intriguingly, Nfu-SatC populated Y-linked region of differentiation in *N. furzeri* and *N. kadleci* (Southern clade) and then, apparently independently, a neo-Y chromosome of *N. brieni*. This result, however, does not tell us whether sex chromosomes of *N. brieni* involve the same synteny block representing ancestral sex chromosomes or whether these systems evolved from entirely different linkage groups. It has been shown that a set of closely related species with demonstrably homeologous sex chromosomes may be populated stochastically by different repetitive sequences such as in Neotropical Triportheidae family [Yano et al., 2016, 2021; Kretschmer et al., 2022]. Inversely, the same or very similar repeat may populate sex chromosomes which evolved independently from different linkage groups [Crepaldi and Parise-Maltempi, 2020; Kretschmer et al., 2022].

C-banding in species with multiple X_1_X_2_Y sex chromosome system revealed moderate degree of constitutive heterochromatin accumulation on neo-Y of *N. brieni* and its low accumulationin *N. ditte* and *N. lourensi.* The latter pattern has been also found on neo-Y of *N. guentheri* [Voleníková et al., 2023]. There findings together suggest that low genetic differentiation is a common feature of *Nothobranchius* X_1_X_2_Y sex chromosome systems, which corroborates the general trend observed among other known fish cases [Sember et al., 2021]. In *N. ditte*, C-banding patterns could not distinguish Y chromosome from the complement, similarly as in *F. thierryi* [Voleníková et al., 2023]. In *N. lourensi*, remarkably, two interstitial heterochromatin blocks were present on its q-arms. These sites may represent regions of former chromosome fusion. Given these patterns together with unusually large Y chromosome size in this species, it is conceivable that this chromosome might have evolved from the fusion of more than two chromosomes during the evolution. Based on the C-banding and FISH patterns, we may, with a high probability, reject the alternative possibility that the Y chromosome size would be enlarged by a massive heterochromatin amplification as reported in other teleosts [Nanda et al., 2014; Schartl et al., 2016]. Fusions of more synteny blocks on the Y chromosome in *N. lourensi* might have led to stronger recombination suppression on this chromosome (particularly around the fusion junctions; cf. [Vara et al., 2021; Yoshida et al., 2023; MacLeod-Bigley and Boulding, 2023]) and this way to facilitate evolution of this sex chromosome. A distinct C-band indicating a probable breakpoint junction has been also revealed on neo-Y of *N. guentheri* [Voleníková et al., 2023].

FISH with PNA telomere probe did not reveal any extra sites in addition to standard location of natural telomeres of all chromosomes. This pattern was observed invariably in all tested individuals across populations and regardless of sex and is consistent also with patterns in *N. furzeri* and *N. kadleci* [Štundlová et al., 2022]. This is intriguing given the obvious high dynamics of *Nothobranchius* spp. karyotype repatterning [Krysanov and Demidova, 2018]. This absence of interstitial telomeric sites (ITSs) is striking especially in species with highly reduced 2n compared to modal chromosome counts (2n = 36–38; [Krysanov and Demidova, 2018]), such as *N. rachovii* (2n = 16) or *N. krysanovi* (2n = 18), where multiple chromosome fusions apparently took place during the evolution. ITSs which may represent remnants of previous rearrangements [Meyne et al., 1990; Ocalewicz, 2013; Vicari et al., 2022] were lacking also on neo-Y chromosomes of *F. thierryi* and four *Nothobranchius* species with X_1_X_2_Y sex chromosome system. As discused above, these Y chromosomes highly likely emerged after fusion of ancestral Y with an autosome. The only cases of ITSs in *Nothobranchius* spp. have been occasionally observed among sampled cells/passages of *N. furzeri* cell lines [Součková et al., in rev.]. The knowledge about the absence of ITS in standard karyotypes at least in the studied species and populations [Štundlová et al., 2022; present study] may be valuable for studies dealing with telomere length meassurements in *Nothobranchius* spp. (e.g., [Reichard et al., 2022a]) by showing that no false positives should be expected to affect the results.

Chromosome fusions and translocations may result i) from telomere attrition or ii) inactivation of telomere or telomere-associated proteins, iii) after double-stranded breaks in centromeric satellite DNA in two monoarmed elements or iv) due to ectopic recombination between abundant repeats on different chromosomes [Slijepcevic, 1996; Schubert and Lysak, 2011; Bolzán, 2012; Stroik and Hendrickson, 2020]. The absence of detectable telomeric repeats may therefore mean their i) complete loss prior/during the rearrangement, ii) too low copy number to be detected by FISH and/or iii) gradual sequence erosion by mutations during evolution [Slijepcevic, 1998; Ocalewicz, 2013]. In favor of the second possibility, telomeric FISH did not yield signals even on natural chromosomal ends of large metacentric pair in *N. cardinalis*, regardless the exposition time. Similar situation has been recently recorded also in Neotropical siluriform fish *Rineloricaria teffeana* [Marajó et al., 2023].

The absence of detectable telomeric repeats in expected/supposed fusion points across all herein studied *Nothobranchius* species suggests that unlike some other fish groups (e.g., [Sember et al., 2015; Marajó et al., 2023]) chromosome rearrangements in *Nothobranchius* seem to follow generally similar mechanistic trajectories and may be driven by similar causes. Fusions of chromosomes with broken or too short telomeres may result from various aberrant events such as replication fork stalling/collapse or telomere uncapping (i.e. loss of associated protective proteins) [Lazzerini-Denchi and Sfeir, 2016; Stroik and Hendrickson, 2020]. We propose that demands for fast lifecycle including rapid growth rate acompanied by massive cell proliferation [Blažek et al., 2013; Cellerino et al., 2016; Vrtílek et al., 2018] might increase the incidence of such events, thereby contributing to the emergence of chromosome rearrangements in *Nothobranchius*. Also prolonged mitotic arrest during diapause [Furness, 2016; Hu and Brunet, 2018] may result in telomere uncapping and subsequent chromosome fusions [Lazzerini-Denchi and Sfeir, 2016]. The resulting chromosome rearrengements might then be faster fixed in small fragmented *Nothobranchius* populations with contribution of genetic drift [King, 1993; Cui et al., 2019] and may be also a subject to meiotic drive process [Voleníková et al., 2023].

## Conclusions

The chromosomal evolution of *Nothobranchius* spp. has been shaped by numerous interchromosomal rearrangements giving rise to a complex karyotype variability, including the emergence of different male-heterogametic sex chromosome systems. By analyzing repetitive DNA patterns on chromosomes of 15 species from four *Nothobranchius* evolutionary lineages, and with the inclusion of outgroup *F. thierryi*, we demonstrated that i) the species share general properties that underpin interchromosomal rearrangements and are manifested by a great reduction or entire elimination of telomeric repeats in the regions of rearrangement; ii) *Nothobranchius* species share the presence of conserved satDNA class (Nfu-SatC) which is thus at least 15 MY old; iii) the species display phylogenetically unrelated variability in Nfu-SatC chromosomal distribution and organization, with occasional occurrence of inter-individual sex-unrelated polymorphisms; iv) Nfu-SatC is not a universal marker for sex chromosome identification as it was found greatly enriched only on the Y chromosome of *N. brieni* in addition to distantly related *N. furzeri* and *N. kadleci* reported previously [Štundlová et al., 2022]. Our data suggests independent trajectories of *Nothobranchius* sex chromosome diferentiation, with recurrent and convergent accumulation of Nfu-SatC on presumably non-recombining Y-linked regions in three species; the remaining representatives with multiple sex chromosomes show generally low degree of differentiation of these systems at cytological level. These findings collectively reinforce the initial presumption of Krysanov and Demidova [2018] that *Nothobranchius* sex chromosomes evolved independently in different species. Repetitive DNA and constitutive heterochromatin patterns revealed in the present study and in Štundlová et al. [2022] and Voleníková et al. [2023] may assist in future genome sequencing attempts in the way to predict potential problems with genome assembly (e.g. [Peona et al., 2020]) and help to propose suitable species and strategy for genome sequencing.

## Supporting information

SuSupporting material for Luksikova et al.

## Acknowledgements

We gratefully acknowledge B. Nagy, A. Nikiforov and H. Hengstler for providing part of the study material and A. Nikiforov for his help in breeding and keeping fishes. We are also grateful to P. Šejnohová and M. Králová for their laboratory assistance, and A. Voleníková (all Institute of Animal Physiology and Genetics, Czech Academy of Sciences, Liběchov, Czech Republic) for repetitive DNA sequence analysis.

## Statement of Ethics

The experimental part involving fish was supervised by the Institutional Animal Care and Use Committee of the Institute of Animal Physiology and Genetics CAS, v.v.i., with the supervisor’s permit number CZ 02361 certified and issued by the Ministry of Agriculture of the Czech Republic. Handling of fish individuals to obtain chromosomes followed European standards in agreement with §17 of the Act No. 246/1992 coll. The experiments with *N. brieni*, *N. ditte*, *N. cardinalis*, *N. foerschi* and one population of *N. guentheri* (NGU2) were approved by the Ethics Committee of Severtsov Institute of Ecology and Evolution (Order No. 27 of November 9, 2018). For direct preparations of chromosomes from the kidney, fish were euthanized using 2-phenoxyethanol (Sigma-Aldrich, St. Louis, MO, USA) before organ sampling. A narrow stripe of the tail fin was taken from live specimens after fish were anaesthetized using MS-222 (Merck KGaA, Darmstadt, Germany).

## Author Contributions

Conceptualization: A. S., M. A., M. R. Data collection: K. L., T. P., M. A., J. Š., Š. P., S. A. S., E. Yu. K., M. J., M. H., A. S. Data analysis: K. L., T. P., M. A., A. S. Writing and original draft preparation: A. S. Writing, review and editing: A. S., P. R., J. Š., S. A. S., M. A., M. R., K. L., T. P., E. Yu. K., M. J.

## Data Availability Statement

All data generated and analysed during this study are within the article and its supplementary material. Further enquiries can be directed to the corresponding author.

## Supporting Information

**Supplementary Table 1.** List of *Nothobranchius* killifish species used in this study, with assignment to their phylogeographic lineage, population/collection codes, source/geographic origin and GPS coordinates of sampling localities.

**Supplementary Figure 2.** Partial karyotypes of *Nothobranchius orthonotus* individuals after single-colour FISH with Nfu-SatC probe (red signals).

**Supplementary Figure 1.** Mitotic metaphases of three *Nothobranchius* species with X_1_X_2_Y sex chromosome system after FISH with Nfu-SatC satDNA and/or C-banding.

## References

Balachandran P, Walawalkar IA, Flores JI, Dayton JN, Audano PA, Beck CR. Transposable element-mediated rearrangements are prevalent in human genomes. Nat Commun. 2022;13(1):7115.

Bartáková V, Reichard M, Blažek R, Polačik M, Bryja J. Terrestrial fishes: Rivers are barriers to gene flow in annual fishes from the African savanna. J Biogeogr. 2015;42(10):1832– 44.

Bartáková V, Reichard M, Janko K, Polačik M, Blažek R, Reichwald K, et al. Strong population genetic structuring in an annual fish, *Nothobranchius furzeri*, suggests multiple savannah refugia in southern Mozambique. BMC Evol Biol. 2013;13:196.

Bellafronte E, Margarido VP, Moreira-Filho O. Cytotaxonomy of *Parodon nasus* and *Parodon tortuosus* (Pisces, Characiformes). A case of synonymy confirmed by cytogenetic analyses. Genet Mol Biol. 2005;28(4):710–6.

Berois N, García G, de Sá RO, editors. Annual fishes: life history strategy, diversity and evolution. Boca Raton, FL: CRC Press; 2016.

Bertollo LAC, Cioffi MB, Moreira-Filho O (2015) Direct chromosome preparation from freshwater teleost fishes. In: Ozouf-Costaz C, Pisano E, Foresti F, and de Almeida-Toledo LF (eds) Fish cytogenetic techniques ray-fin fishes and chondrichthyans. CRC Press, Inc, Endfield, pp 21–26. https://doi.org/10.1201/b18534-4

Biscotti MA, Canapa A, Forconi M, Olmo E, Barucca M. Transcription of tandemly repetitive DNA: functional roles. Chromosom Res. 2015;23(3):463–77.

Blažek R, Polačik M, Kačer P, Cellerino A, Řežucha R, Methling C, et al. Repeated intraspecific divergence in life span and aging of African annual fishes along an aridity gradient. Evolution. 2017;71(2):386–402.

Blažek R, Polačik M, Reichard M. Rapid growth, early maturation and short generation time in African annual fishes. Evodevo. 2013;4(1):24.

Blommaert J, Riss S, Hecox-Lea B, Mark Welch DB, Stelzer CP. Small, but surprisingly repetitive genomes: Transposon expansion and not polyploidy has driven a doubling in genome size in a metazoan species complex. BMC Genomics. 2019;20(1):466.

Bolzán AD. Chromosomal aberrations involving telomeres and interstitial telomeric sequences. Mutagenesis. 2012;27(1):1–15.

Bracewell R, Chatla K, Nalley MJ, Bachtrog D. Dynamic turnover of centromeres drives karyotype evolution in *Drosophila*. Elife. 2019;8:1–47.

Brown RE, Freudenreich CH. Structure-forming repeats and their impact on genome stability. Curr Opin Genet Dev. 2021;67:41–51.

Camacho JPM, Cabrero J, López-León MD, Martín-Peciña M, Perfectti F, Garrido-Ramos MA, et al. Satellitome comparison of two oedipodine grasshoppers highlights the contingent nature of satellite DNA evolution. BMC Biol. 2022;20(1):36.

Cellerino A, Valenzano DR, Reichard M. From the bush to the bench: The annual *Nothobranchius* fishes as a new model system in biology. Biol Rev Camb Philos Soc. 2016;91(2):511–33.

Cioffi MB, Bertollo LAC. Chromosomal distribution and evolution of repetitive DNAs in fish. Genome Dyn. 2012;7:197–221.

Crepaldi C, Parise-Maltempi PP. Heteromorphic sex chromosomes and their DNA content in fish: An insight through satellite DNA accumulation in *Megaleporinus elongatus*. Cytogenet Genome Res. 2020;160(1):38–46.

Cui R, Medeiros T, Willemsen D, Iasi LNM, Collier GE, Graef M, et al. Relaxed selection limits lifespan by increasing mutation load. Cell. 2019;178(2):385–399.e20.

Deon GA, Glugoski L, Vicari MR, Nogaroto V, Sassi F de MC, Cioffi MB, et al. Highly rearranged karyotypes and multiple sex chromosome systems in armored catfishes from the genus *Harttia* (Teleostei, Siluriformes). Genes (Basel). 2020;11(11):1–17.

Dorn A, Musilová Z, Platzer M, Reichwald K, Cellerino A. The strange case of East African annual fish: Aridification correlates with diversification for a savannah aquatic group? BMC Evol Biol. 2014;14(1):210.

Dos Santos RZ, Calegari RM, Silva DMZ de A, Ruiz-Ruano FJ, Melo S, Oliveira C, et al. A long-term conserved satellite DNA that remains unexpanded in several genomes of Characiformes fish is actively transcribed. Genome Biol Evol. 2021;13(2):evab002.

Ewulonu UV, Haas R, Turner BJ. A multiple sex chromosome system in the annual killifish, *Nothobranchius guentheri*. Copeia. 1985;1985(2):503–8.

Feliciello I, Akrap I, Brajkovi J, Zlatar I, Ugarkovic D. Satellite DNA as a driver of population divergence in the red flour beetle *Tribolium castaneum*. Genome Biol Evol. 2014;7(1):228–39.

Feliciello I, Pezer Ž, Sermek A, Mađarić BB, Ljubić S, Ugarković Đ. 2021 Satellite DNA-mediated gene expression regulation: Physiological and evolutionary implication. Ð. Ugarković (ed.), Satellite DNAs in Physiology and Evolution, Progress in Molecular and Subcellular Biology 60, https://doi.org/10.1007/978-3-030-74889-0_6

Ferree PM, Barbash DA. Species-specific heterochromatin prevents mitotic chromosome segregation to cause hybrid lethality in *Drosophila*. PLoS Biol. 2009;7(10):e1000234.

Fricke R, Eschmeyer WN, Van der Laan R (eds) (2022) Eschmeyer’s catalog of fishes: genera, species, references. http://researcharchive.calacademy.org/research/ichthyology/catalog/fishcatmain.asp. Accessed 10 January 2023

Fry K, Salser W. Nucleotide sequences of HS-α satellite DNA from kangaroo rat *Dipodomys ordii* and characterization of similar sequences in other rodents. Cell. 1977;12(4):1069– 84.

Furness AI. The evolution of an annual life cycle in killifish: Adaptation to ephemeral aquatic environments through embryonic diapause. Biol Rev Camb Philos Soc. 2016;91:796– 812.

Garrido-Ramos MA. Satellite DNA: An evolving topic. Genes (Basel). 2017;8(9):230.

Glugoski L, Deon G, Schott S, Vicari MR, Nogaroto V, Moreira-Filho O. Comparative cytogenetic analyses in *Ancistrus species* (Siluriformes: Loricariidae). Neotrop Ichthyol. 2020;18(2):e200013.

Goes CAG, Daniel SN, Piva LH, Yasui GS, Artoni RF, Hashimoto DT, et al. Cytogenetic markers as a tool for characterization of hybrids of *Astyanax* Baird & Girard, 1854 and *Hyphessobrycon* Eigenmann, 1907. Comp Cytogenet. 2020;14(2):231–42.

Goes CAG, Dos Santos N, Rodrigues PHM, Stornioli JHF, Silva ABD, Dos Santos RZ, et al. The satellite DNA catalogues of two Serrasalmidae (Teleostei, Characiformes): Conservation of general satDNA features over 30 million years. Genes (Basel). 2022;14(1):91.

Haaf T, Schmid M. An early stage of ZZ/ZW sex chromosomes differentiation in *Poecilia sphenops* var. *melanistica* (Poeciliidae, Cyprinodontiformes). Chromosoma. 1984;89:37– 41.

Hartley G, O’Neill RJ. Centromere repeats: Hidden gems of the genome. Genes (Basel). 2019;10(3):223.

Hu CK, Brunet A. The African turquoise killifish: A research organism to study vertebrate aging and diapause. Aging Cell. 2018;17(3):e12757.

Charlesworth B, Sniegowski P, Stephan W. The evolutionary dynamics of repetitive DNA in eukaryotes. Nature. 1994;371(6494):215–20.

Charlesworth D. Young sex chromosomes in plants and animals. New Phytol. 2019;224(3):1095–107.

Kent TV, Uzunović J, Wright SI. Coevolution between transposable elements and recombination. Philos Trans R Soc Lond B Biol Sci. 2017;372(1736):20160458.

King M (1993) Species Evolution: The Role of Chromosomal Change. Cambridge University Press, Cambridge, UK.

Kligerman AD, Bloom SE. Rapid chromosome preparations from solid tissues of fishes. J Fish Res Board Can. 1977;34(2):266–9.

Kretschmer R, Goes CAG, Bertollo LAC, Ezaz T, Porto-Foresti F, Toma GA, et al. Satellitome analysis illuminates the evolution of ZW sex chromosomes of Triportheidae fishes (Teleostei: Characiformes). Chromosoma. 2022;131(1–2):29–45.

Krysanov E, Demidova T, Nagy B. Divergent karyotypes of the annual killifish genus *Nothobranchius* (Cyprinodontiformes, Nothobranchiidae). Comp Cytogenet. 2016;10(3):439–45.

Krysanov E, Demidova T. Extensive karyotype variability of African fish genus *Nothobranchius* (Cyprinodontiformes). Comp Cytogenet. 2018;12(3):387–402.

Krysanov EY, Nagy B, Watters BR, Sember A, Simanovsky SA. Karyotype differentiation in the *Nothobranchius ugandensis* species group (Teleostei, Cyprinodontiformes), seasonal fishes from the east African inland plateau, in the context of phylogeny and biogeography. Comp Cytogenet. 2023;7(1):13–29.

Lazzerini-Denchi E, Sfeir A. Stop pulling my strings – what telomeres taught us about the DNA damage response. Nat Rev Mol Cell Biol. 2016;17(6):364–78.

Lee J, Waminal NE, Choi H Il, Perumal S, Lee SC, Nguyen VB, et al. Rapid amplification of four retrotransposon families promoted speciation and genome size expansion in the genus *Panax*. Sci Rep. 2017;7(1):9045.

Levan AK, Fredga K, Sandberg AA. Nomenclature for centromeric position on chromosomes. Hereditas. 1964;52(2):201–20.

Li SF, Su T, Cheng GQ, Wang BX, Li X, Deng CL, et al. Chromosome evolution in connection with repetitive sequences and epigenetics in plants. Genes (Basel). 2017;8(10):290.

Lichilín N, Salzburger W, Böhne A. No evidence for sex chromosomes in natural populations of the cichlid fish *Astatotilapia burtoni*. G3 (Bethesda). 2023. https://doi.org/10.1093/g3journal/jkad011

López-Flores I, Garrido-Ramos MA. The repetitive DNA content of eukaryotic genomes. Genome Dyn. 2012;7:1–28.

Louzada S, Lopes M, Ferreira D, Adega F, Escudeiro A, Gama-Carvalho M, et al. Decoding the role of satellite DNA in genome architecture and plasticity – an evolutionary and clinical affair. Genes. 2020;11(1):72.

Lower SS, McGurk MP, Clark AG, Barbash DA. Satellite DNA evolution: Old ideas, new approaches. Curr Opin Genet Dev. 2018;49:70–8.

MacPherson N, Champion CP, Weir LK, Dalziel AC. Reproductive isolating mechanisms contributing to asymmetric hybridization in Killifishes (*Fundulus* spp.). J Evol Biol. 2023. https://doi.org/10.1111/jeb.14148

Marajó L, Viana PF, Ferreira AMV, Py-Daniel LHR, Cioffi MB, Sember A, et al. Chromosomal rearrangements and the first indication of an ♀X_1_X_1_X_2_X_2_/♂ X_1_X_2_Y sex chromosome system in *Rineloricaria* fishes (Teleostei: Siluriformes). J Fish Biol. 2023;102(2), 443–454. https://doi.org/10.1111/jfb.15275

Marta A, Dedukh D, Bartoš O, Majtanová Z, Janko K. Cytogenetic characterization of seven novel satDNA markers in two species of spined loaches (*Cobitis*) and their clonal hybrids. Genes. 2020;11(6):617.

Melters DP, Bradnam KR, Young HA, Telis N, May MR, Ruby JG, et al. Comparative analysis of tandem repeats from hundreds of species reveals unique insights into centromere evolution. Genome Biol. 2013;14(1):R10.

Meštrović N, Mravinac B, Pavlek M, Vojvoda-Zeljko T, Šatović E, Plohl M. Structural and functional liaisons between transposable elements and satellite DNAs. Chromosome Res. 2015;23(3):583–96.

Meyne J, Baker RJ, Hobart HH, Hsu TC, Ryder OA, Ward OG, et al. Distribution of non-telomeric sites of the (TTAGGG)n telomeric sequence in vertebrate chromosomes. Chromosoma. 1990;99(1):3–10.

Nagy B. *Nothobranchius ditte*, a new species of annual killifish from the Lake Mweru basin in the Democratic Republic of the Congo (Teleostei: Nothobranchiidae). Ichthyo-logical Exploration of Freshwaters 2018;28(2):115–134.

Nagy B, Watters BR. A review of the conservation status of seasonal *Nothobranchius* fishes (Teleostei: Cyprinodontiformes), a genus with a high level of threat, inhabiting ephemeral wetland habitats in Africa. Aquat Conserv: Mar Freshw Ecosyst. 2022;32(1):199–216.

Nanda I, Schories S, Tripathi N, Dreyer C, Haaf T, Schmid M, et al. Sex chromosome polymorphism in guppies. Chromosoma. 2014;123(4):373–83.

Nonaka E, Sirén J, Somervuo P, Ruokolainen L, Ovaskainen O, Hanski I. Scaling up the effects of inbreeding depression from individuals to metapopulations. J Anim Ecol. 2019;88(8):1202–14.

Novák P, Neumann P, Macas J. Global analysis of repetitive DNA from unassembled sequence reads using RepeatExplorer2. Nat Protoc. 2020;15(11):3745–76.

O’Sullivan RJ, Karlseder J. Telomeres: protecting chromosomes against genome instability. Nat Rev Mol Cell Biol. 2010;11(3):171–81.

Ocalewicz K. Telomeres in fishes. Cytogenet Genome Res. 2013;141(2–3):114–25.

Peona V, Blom MPK, Xu L, Burri R, Sullivan S, Bunikis I, et al. Identifying the causes and consequences of assembly gaps using a multiplatform genome assembly of a bird-of-paradise. Mol Ecol Resour. 2020;21:263–86.

Plohl M, Meštrović N, Mravinac B. Satellite DNA evolution. Genome Dyn. 2012;7:126–52.

Polačik M, Reichard M. Asymmetric reproductive isolation between two sympatric annual killifish with extremely short lifespans. PLoS One. 2011;6(8):e22684.

Ráb P, Roth P (1988) Cold-blooded vertebrates. In: Balicek P, Forejt J, Rubeš J (eds) Methods of chromosome analysis. Cytogenetická Sekce Československé Biologické Společnosti při CSAV, Brno, Czech Republic, pp 115–124

Reichard M, Giannetti K, Ferreira T, Maouche A, Vrtílek M, Polačik M, et al. Lifespan and telomere length variation across populations of wild-derived African killifish. Mol Ecol. 2022a;31(23):5979–92.

Reichard M, Janáč M, Blažek R, Žák J, Alila OD, Polačik M. Patterns and drivers of *Nothobranchius* killifish diversity in lowland Tanzania. Ecol Evol. 2022b;12(6):e8990.

Reichwald K, Lauber C, Nanda I, Kirschner J, Hartmann N, Schories S, et al. High tandem repeat content in the genome of the short-lived annual fish *Nothobranchius furzeri*: A new vertebrate model for aging research. Genome Biol. 2009;10(2):R16.

Reichwald K, Petzold A, Koch P, Downie BR, Hartmann N, Pietsch S, et al. Insights into sex chromosome evolution and aging from the genome of a short-lived fish. Cell. 2015;163(6):1527–38.

Robledillo LA, Neumann P, Koblížková A, Novák P, Vrbová I, Macas J. Extraordinary sequence diversity and promiscuity of centromeric satellites in the legume Tribe Fabeae. Mol Biol Evol. 2020;37(8):2341–56.

Robles F, De La Herrán R, Ludwig A, Ruiz Rejón C, Ruiz Rejón M, Garrido-Ramos MA. Evolution of ancient satellite DNAs in sturgeon genomes. Gene. 2004;338(1):133–42.

Ruiz-Ruano FJ, López-León MD, Cabrero J, Camacho JPM. High-throughput analysis of the satellitome illuminates satellite DNA evolution. Sci Rep. 2016;6(1):28333.

Sassi F de MC, Deon GA, Moreira-Filho O, Vicari MR, Bertollo LAC, Liehr T, et al. Multiple sex chromosomes and evolutionary relationships in amazonian catfishes: The outstanding model of the genus *Harttia* (Siluriformes: Loricariidae). Genes (Basel). 2020;11(10):1– 16.

Sember A, Bohlen J, Šlechtová V, Altmanová M, Symonová R, Ráb P. Karyotype differentiation in 19 species of river loach fishes (Nemacheilidae, Teleostei): Extensive variability associated with rDNA and heterochromatin distribution and its phylogenetic and ecological interpretation. BMC Evol Biol. 2015;15:251.

Sember A, Nguyen P, Perez MF, Altmanová M, Ráb P, Cioffi MB. Multiple sex chromosomes in teleost fishes from a cytogenetic perspective: state of the art and future challenges. Phil Trans R Soc B. 2021;376(1833):20200098.

Serrano-Freitas ÉA, Silva DMZA, Ruiz-Ruano FJ, Utsunomia R, Araya-Jaime C, Oliveira C, et al. Satellite DNA content of B chromosomes in the characid fish *Characidium gomesi* supports their origin from sex chromosomes. Mol Genet Genomics. 2020;295(1):195– 207.

Shen G, Wang P (2019) Environmental sex determination and sex differentiation in teleosts – how sex is established. In: (eds. Wang HP, Piferrer F, Chen S-L (eds) Sex Control in Aquaculture, 1st edn. John Wiley & Sons, Hoboken, pp 85–115.

Schartl M, Georges A, Graves JAM. Polygenic sex determination in vertebrates – is there any such thing? Trends Genet. 2023.

Shao C, Sun S, Liu K, Wang J, Li S, Liu Q, Deagle BE, Seim I, Biscontin A, Wang Q, et al. The enormous repetitive Antarctic krill genome reveals environmental adaptations and population insights. Cell. 2023;186:1–16.

Schemberger MO, Bellafronte E, Nogaroto V, Almeida MC, Schühli GS, Artoni RF, et al. Differentiation of repetitive DNA sites and sex chromosome systems reveal closely related group in Parodontidae (Actinopterygii: Characiformes). Genetica. 2011;139(11– 12):1499–508.

Schemberger MO, Nascimento VD, Coan R, Ramos É, Nogaroto V, Ziemniczak K, et al. DNA transposon invasion and microsatellite accumulation guide W chromosome differentiation in a Neotropical fish genome. Chromosoma. 2019;128(4):547–60.

Schubert I, Lysák MA. Interpretation of karyotype evolution should consider chromosome structural constraints. Trends Genet. 2011;27(6):207–16.

Slijepcevic P. Telomeres and mechanisms of Robertsonian fusion. Chromosoma. 1998;107(2):136–40.

Sochorová J, Garcia S, Gálvez F, Symonová R, Kovařík A. Evolutionary trends in animal ribosomal DNA loci: Introduction to a new online database. Chromosoma. 2018;127(1):141–50.

Součková K, Jasík M, Sovadinová I, Sember A, Sychrová E, Konieczna A, Bystrý V, Dyková I, Blažek R, Lukšíková K, et al. From fish to cells: Establishment of continuous cell lines from embryos of annual killifish *Nothobranchius furzeri* and *N. kadleci*. Aquat Toxicol. under review

Stephan W. Recombination and the evolution of satellite DNA. Genet Res. 1986;47(3):167–74.

Stroik S, Hendrickson EA. Telomere fusions and translocations: A bridge too far? Curr Opin Genet Dev. 2020;60:85–91.

Šatović-Vukšić E, & Plohl M. Satellite DNAs—from localized to highly dispersed genome components. Genes. 2023;14(3):742.

Štundlová J, Hospodářská M, Lukšíková K, Voleníková A, Pavlica T, Altmanová M, et al. Sex chromosome differentiation via changes in the Y chromosome repeat landscape in African annual killifishes *Nothobranchius furzeri* and *N. kadleci*. Chromosome Res. 2022;30(4):309–33.

Talbert PB, Henikoff S. What makes a centromere? Exp Cell Res. 2020;389(2):111895.

Thakur J, Packiaraj J, Henikoff S. Sequence, chromatin and evolution of satellite DNA. Int J Mol Sci. 2021;22(9):4309.

Underwood CJ, Choi K. Heterogeneous transposable elements as silencers, enhancers and targets of meiotic recombination. Chromosoma. 2019;128(3):279–96.

Utsunomia R, Silva DMZ de A, Ruiz-Ruano FJ, Goes CAG, Melo S, Ramos LP, et al. Satellitome landscape analysis of *Megaleporinus macrocephalus* (Teleostei, Anostomidae) reveals intense accumulation of satellite sequences on the heteromorphic sex chromosome. Sci Rep. 2019;9(1):5856.

van der Merwe PDW, Cotterill FPD, Kandziora M, Watters BR, Nagy B, Genade T, et al. Genomic fingerprints of palaeogeographic history: The tempo and mode of rift tectonics across tropical Africa has shaped the diversification of the killifish genus *Nothobranchius* (Teleostei: Cyprinodontiformes). Mol Phylogenet Evol. 2021;158:106988.

Vara C, Paytuví-Gallart A, Cuartero Y, Álvarez-González L, Marín-Gual L, Garcia F, et al. The impact of chromosomal fusions on 3D genome folding and recombination in the germ line. Nat Commun. 2021;12(1):2981.

Vicari MR, Bruschi DP, Cabral-de-Mello DC. Telomere organization and the interstitial telomeric sites involvement in insects and vertebrates chromosome evolution. Genet Mol Biol. 2022;45(3 Suppl 1):e20220071.

Vicoso B. Molecular and evolutionary dynamics of animal sex-chromosome turnover. Nat Ecol Evol. 2019;3(12):1632–41.

Voleníková A, Lukšíková K, Mora P, Pavlica T, Altmanová M, Štundlová J, Pelikánová Š, Simanovsky SA, Jankásek M, Reichard M, Nguyen P, Sember A. Fast centromeric repeat turnover provides a glimpse into satellite DNA evolution in *Nothobranchius* annual killifishes. bioRxiv [Preprint]. 2023. doi: 10.1101/2023.03.23.534043.

Völker M, Ráb P (2015) Direct chromosome preparation from regenerating fin tissue. In: Ozouf-Costaz C, Pisano E, Foresti F, and de Almeida-Toledo LF (eds) Fish cytogenetic techniques: ray-fin fishes and chondrichthyans. CRC Press, Inc, Endfield, pp 37–41. https://doi.org/10.1201/b18534-4

Vondrak T, Robledillo LA, Novák P, Koblížková A, Neumann P, Macas J. Characterization of repeat arrays in ultra-long nanopore reads reveals frequent origin of satellite DNA from retrotransposon-derived tandem repeats. Plant J. 2020;101(2):484–500.

Vrtílek M, Žák J, Pšenička M, Reichard M. Extremely rapid maturation of a wild African annual fish. Curr Biol. 2018;28(15):R822–4.

Watters, B.W., B.J. Cooper and R.H. Wildekamp. Description of *Nothobranchius cardinalis* spec. nov. (Cyprinodontiformes: Aplocheilidae), an annual fish from the Mbwemkuru River basin, Tanzania. J. Am. Killifsh Ass. 2008;40(5&6):129–145.

Watters, B. R., B. Nagy, P. D. W. van der Merwe, F. P. D. Cotterill & D. U. Bellstedt. 2020. Redescription of the seasonal killifish species *Nothobranchius ocellatus* and description of a related new species *Nothobranchius matanduensis*, from eastern Tanzania (Teleostei: Nothobranchiidae). Ichthyol Explor Freshw. 2020;30 (2):151–178.

Wildekamp RH (1996) A world of killies. Atlas of the oviparous cyprinodontiform fishes of the world (Vol. III). American Killifish Association, Mishawaka, 330 pp.

Wildekamp RH (2004) A world of killies – atlas of the oviparous cyprinodontiform fishes of the world (Vol. IV). The American Killifish Association, Elyria, Ohio, 398 pp.

Willemsen D, Cui R, Reichard M, Valenzano DR. Intra-species differences in population size shape life history and genome evolution. Elife. 2020;9:e55794.

Winter DJ, Ganley ARD, Young CA, Liachko I, Schardl CL, Dupont PY, et al. Repeat elements organize 3D genome structure and mediate transcription in the filamentous fungus *Epichloë festucae*. PLoS Genet. 2018;14(10):e1007467.

Yano CF, Bertollo LAC, Ezaz T, Trifonov V, Sember A, Liehr T, et al. Highly conserved Z and molecularly diverged W chromosomes in the fish genus *Triportheus* (Characiformes, Triportheidae). Heredity (Edinb). 2017;118(3):276–83.

Yano CF, Bertollo LAC, Liehr T, Troy WP, Cioffi MB. W chromosome dynamics in *Triportheus* species (Characiformes, Triportheidae): An ongoing process narrated by repetitive sequences. J Hered. 2016;107(4):342–8.

Yano CF, Sember A, Kretschmer R, Bertollo LAC, Ezaz T, Hatanaka T, et al. Against the mainstream: Exceptional evolutionary stability of ZW sex chromosomes across the fish families Triportheidae and Gasteropelecidae (Teleostei: Characiformes). Chromosome Res. 2021;29(3–4):391–416.

Yoshida K, Rödelsperger C, Röseler W, Riebesell M, Sun S, Kikuchi T, Sommer RJ. Chromosome fusions repatterned recombination rate and facilitated reproductive isolation during *Pristionchus* nematode speciation. Nat. Ecol. Evolut. 2023;7:424–439

Zeljko VT, Pavlek M, Meštrović N, Plohl M. Satellite DNA-like repeats are dispersed throughout the genome of the Pacific oyster *Crassostrea gigas* carried by Helentron non-autonomous mobile elements. Sci Rep. 2020;10(1):15107.

