## Supplementary material for "Conserved satellite DNA motif and lack of interstitial telomeric sites in highly rearranged African *Nothobranchius* killifish karyotypes": SuSupporting material for Luksikova et al.

**The PDF file includes:**

Supplementary Table 1
Supplementary Figure 1
Supplementary Figure 2

**Supplementary Table 1.** List of *Nothobranchius* killifish species used in this study, with assignment to their phylogeographic lineage, population/collection codes, source/geographic origin and GPS coordinates of sampling localities.

| Clade | Species | Population (collection code) | Source/ locality | GPS coordinates |
|---|---|---|---|---|
| outgroup | *Fundulosoma thierryi* Ahl, 1924 | aquarium strain | — | — |
| Southern clade | *Nothobranchius orthonotus* (Peters, 1844) | MZCS-02 | Limpopo (Mozambique) | 24°03'48.5"S 32°43'55.9"E |
| | *N. kuhntae* (Ahl, 1926) | MZCS-528 | Pungwe (Mozambique) | 19°41'50.5"S 34°46'58.6"E |
| | *N. pienaari* Shidlovskyi, Watters, Wildekamp, 2010 | MZCS-505 | Limpopo (Mozambique) | 23°31'47.2"S 32°34'40.6"E |
| | | MZCS-514 | Pungwe (Mozambique) | 19°41'57.1"S 34°47'05.0"E |
| | *N. krysanovi* Shidlovskyi, Watters, Wildekamp, 2010 | aquarium strain; MZCS-249 | Quelimane (Mozambique) | 17°48'52.2"S 36°54'49.4"E |
| | *N. rachovii* Ahl, 1926 | MZCS-096 | Beira Airport (Mozambique) | 19°48'48.8"S 34°54'17.6"E |
| Ocellatus clade | *N. ocellatus* (Seegers, 1985) | — | Nyamwage (Tanzania) | — |
| Coastal clade | *N. eggersi* Seegers, 1982 | T52 | Bagamoyo (Tanzania) | 6°28'55.9"S 38°54'51.5"E |
| | *N. lourensi* Wildekamp, 1977 | TZH 2018-101 | Ifakara (Tanzania) | 8°10'00.0"S 36°41'30.0"E |
| | *N. foerschi* Wildekamp, Berkenkamp, 1979 | CI 57 | Soga (Tanzania) | 6°50'13.2"S 38°50'45.6"E |
| | *N. guentheri* (Pfeffer, 1983) | aquarium strain | Zanzibar (Tanzania) | — |
| | | ZAN 14-02 | Zanzibar (Tanzania) | 5°58'43.2"S 39°14'58.2"E |
| | *N. cardinalis* Watters, Cooper, Wildekamp, 2008 | TTKSN 17-12 | Matandu (Tanzania) | 9°30'04.0"S 38°13'49.0"E |
| | *N. rubripinnis* Seegers, 1986 | T33 | Kitonga (Tanzania) | 7°12'40.5"S 39°10'30.9"E |
| | *N. melanospilus* (Pfeffer, 1986) | T163 | Makurange (Tanzania) | 6°28'02.8"S 38°47'55.3"E |
| | | T123 | Nyamwage (Tanzania) | 8°04'29.2"S 39°00'08.4"E |
| Kalahari clade | *N. brieni* Poll, 1938 | CD 13-4 | Bukama (Kongo) | 9°17'14.0"S 25°56'34.0"E |
| | *N. ditte* Nagy, 2018 | CD 16-13 | Kilwa (Kongo) | 9°12'33.0"S 28°17'01.0"E |

Additional information to the origin of the sampled individuals is provided in the main text, section “Fish Sampling”. Species order reflects their phylogenetic relationships ([van der Merwe et al., 2021] and Figure 1).

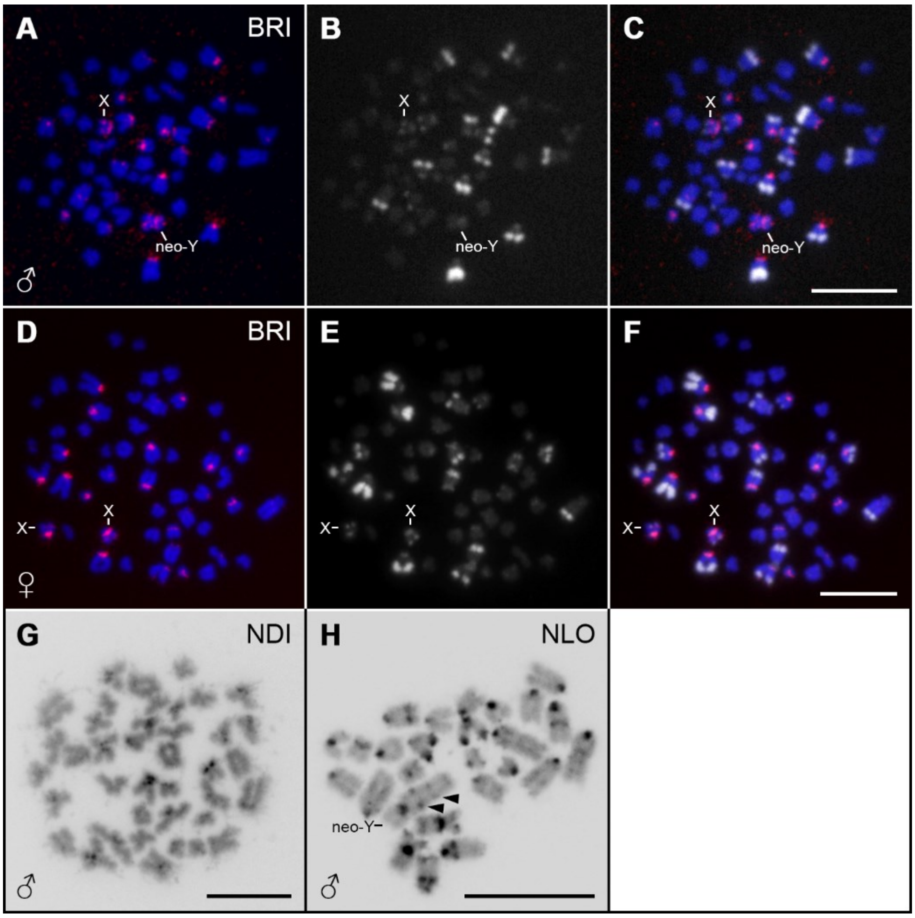


**Supplementary Figure 1.** Mitotic metaphases of three *Nothobranchius* species with $X_1X_2Y$ sex chromosome system after FISH with Nfu-SatC satDNA and/or C-banding. *N. brieni* male (A–C) and *N. brieni* female (D–F) after sequential cytogenetic analysis: A, D) FISH with Nfu-SatC probe (red signals); DAPI-counterstained chromosomes (blue); B, E) C-banding with DAPI counterstaining, performed sequentially on the same metaphases as Nfu-SatC FISH; C, F) merged images of Nfu-SatC and C-banding patterns. G–H) C-banding with DAPI counterstaining (inverted pictures) in G) *N. ditte* and H) *N. lourensi* males. Neo-Y and one of the X chromosomes are depicted if unambiguously detectable. Note the prominent accumulation of Nfu-SatC on the Y chromosome of *N. brieni* (A). Notice also largely non-overlapping patterns of Nfu-SatC distribution and C-banding. Full arrowheads denote interstitial heterochromatin blocks on remarkably large Y chromosome of *N. lourensi* (H). The metaphase spread of *N. lourensi* (H) displays incomplete 2n (26 chromosomes; one chromosome missing), however, it provides the best spreading and sufficient data to present C-banding results (especially regarding pattern details on Y chromosome). Species coding follows Figure 1. Scale bar = 10 μm.

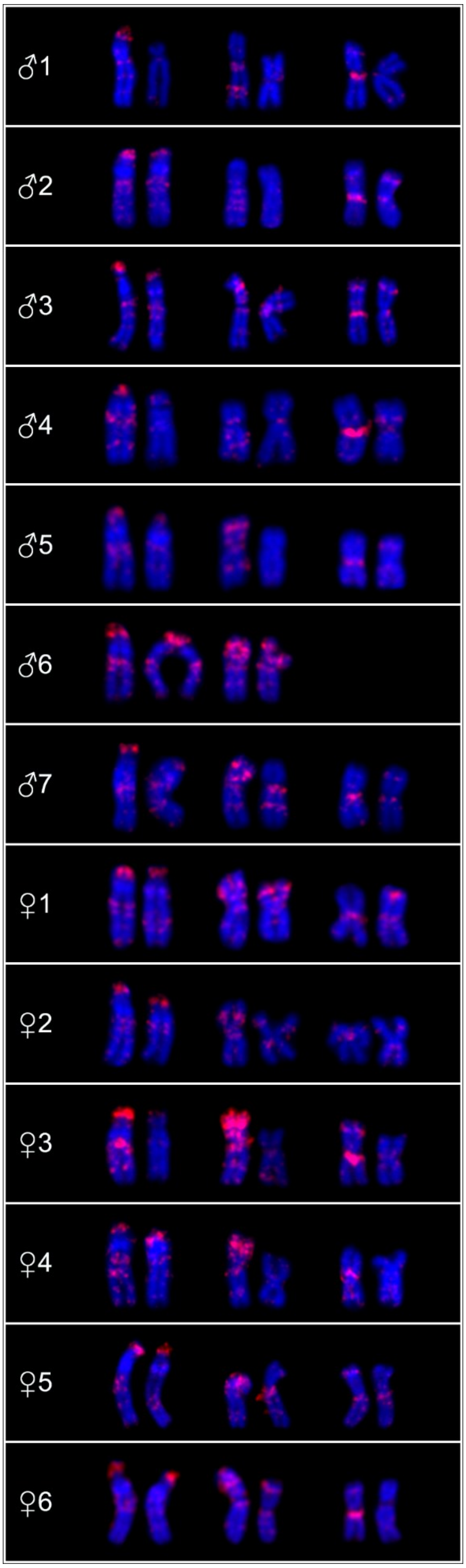


**Supplementary Figure 2.** Partial karyotypes of *Nothobranchius orthonotus* individuals after single-colour FISH with Nfu-SatC probe (red signals). The chromosome pairs with notable polymorphic Nfu-SatC patterns were selected. For further explanations, see main text.
